# Fetal MRI reveals altered prenatal cortical surface area in fetuses later diagnosed with autism spectrum disorder

**DOI:** 10.64898/2026.06.01.729342

**Authors:** Andrea Gondova, Suzette Saucedo Olvera, Yinong Qiu, Haiyi Guo, Seungyoon Jeong, Wendy K. Chung, P. Ellen Grant, Xue-Jun Kong, Kiho Im

## Abstract

Autism spectrum disorder (ASD) is increasingly conceptualized as a condition rooted in altered prenatal neurodevelopment, yet in vivo evidence from fetal brain imaging remains limited. Using retrospective fetal MRI and surface-based morphometry, we investigated cortical development in 15 fetuses later diagnosed with ASD (77%; mean gestational age [GA] = 26.7 weeks) without major structural brain abnormalities and compared them with 60 typically developing controls (57% male; mean GA = 28.4 weeks). Fetuses later diagnosed with ASD showed significantly reduced whole-brain inner cortical plate surface area compared with controls (β = −0.08 ± 0.02 SE, p = 0.002, partial η² = 0.13), corresponding to an estimated ∼7.7% reduction (predicted at GA = 28.1 weeks). Lobar mixed-effects analyses demonstrated broadly distributed reductions across all cortical lobes (FDR-corrected p = <0.001–0.024; Cohen’s d = −0.06 to −0.10), with modest regional heterogeneity indicating relatively greater frontal and insular involvement (group×lobe: F = 19.31, p = 0.002, η² = 0.08). Surface area findings remained directionally stable across sensitivity analyses, including restriction to neurodevelopmentally confirmed controls and models accounting for image quality variability, although effect sizes were attenuated after quality adjustment. Normative modeling further demonstrated subtle negative deviations from typical prenatal cortical surface area trajectories in ASD (mean Z = −0.27, p = 0.018). These findings suggest that aspects of cortical morphogenesis may diverge prenatally in individuals later diagnosed with ASD and suggest the feasibility of fetal MRI-based surface morphometry for studying early neurodevelopmental variation associated with ASD risk.

## 1. Introduction

Autism spectrum disorder (ASD) affects approximately 1-2% of children worldwide, with evidence suggesting a continued increase in prevalence (Shaw et al., 2025). Core features include impairments in social communication and interaction, together with restricted and repetitive behaviors (American Psychiatric Association, 2022). Beyond these features, ASD is frequently accompanied by psychiatric and medical comorbidities, with ∼70% of individuals experiencing at least one co-occurring condition and a more than twofold increased risk of premature mortality (O’Nions et al., 2024), contributing to substantial lifelong burden on individuals, families, and healthcare systems. However, diagnosis currently remains behaviorally defined and typically not reliable until 2-3 years of age, after symptom emergence, limiting opportunities for pre-symptomatic risk stratification and early prevention (Brian et al., 2019). These limitations have motivated extensive work aimed at improving our understanding of early brain differences associated with ASD and the search for early biologically grounded markers using neuroimaging approaches (Clairmont et al., 2022; Halliday et al., 2024; Hiremath et al., 2021).

ASD is increasingly conceptualized as a condition rooted in altered early neurodevelopment (Amgalan et al., 2021), with converging genetic, epidemiological, and neuropathological evidence suggesting that relevant biological processes begin during prenatal life (Motta et al., 2026). Large-scale genomic studies implicate genes involved in neurogenesis, neuronal differentiation, synaptic organization, and cortical development (Gogate et al., 2024), while epidemiological associations with maternal immune, metabolic, and environmental factors further support the importance of the intrauterine environment during critical periods of brain development (Holland et al., 2026; Love et al., 2024). However, most neuroimaging studies focus on postnatal stages (Halliday et al., 2024), when observed differences may reflect a combination of earlier developmental trajectories, postnatal environmental influences, and adaptive or compensatory processes that complicate interpretation (Batalle et al., 2018; Dawson, 2008; Nava & Röder, 2011; Tooley et al., 2021). Earlier developmental windows may thus be particularly informative for understanding the origins of ASD-related neurodevelopmental differences.

Recent advances in fetal magnetic resonance imaging (MRI) enable in vivo characterization of prenatal cortical development (Gholipour et al., 2011; Prayer et al., 2023; Wyburd et al., 2026) and are increasingly sensitive to deviations associated with clinical conditions and in utero exposures (Ahmad et al., 2023; Stuempflen et al., 2023; Wilson et al., 2025). Fetal MRI provides an unique opportunity to study brain development during periods of rapid cortical expansion and organization that establish the structural and functional architecture of emerging neural systems that support optimal outcomes (Gilmore et al., 2018). To date, research linking prenatal neuroimaging to later ASD diagnosis remains limited but is increasingly informative. Structural fetal abnormalities such as ventriculomegaly and subependymal lesion burden in tuberous sclerosis complex have been associated with elevated ASD risk (Hulshof et al., 2021; Kyriakopoulou et al., 2023), and fetal functional MRI studies have suggested that altered in utero connectivity may relate to later autistic traits (Chen et al., 2026). Retrospective fetal imaging studies further support the presence of early neuroanatomical differences, reporting regional alterations in cortical volumetry and hemispheric asymmetry, including increased insular cortex volume in children later diagnosed with ASD (Ortug et al., 2024).

Here, we extend this work by focusing on cortical surface area as a complementary marker of early brain development, motivated by postnatal studies suggesting its relevance to ASD-related cortical differences (Mensen et al., 2017; Ohta et al., 2016). Surface area reflects tangential cortical expansion and sulcation-related developmental processes that increase the spatial availability for white matter connectivity, and is influenced by genetic factors partially distinct from those associated for example with cortical thickness (Panizzon et al., 2009). It may be particularly sensitive to alterations in early neurogenesis, progenitor cell proliferation, and radial unit expansion, providing complementary information to volumetric measures, and potentially serving as an informative marker of atypical prenatal cortical development associated with ASD.

In this study, we investigated prenatal cortical development in a retrospective cohort of fetuses later diagnosed with ASD. While major structural brain malformations are expected to confer elevated risk for later ASD and represent an important context for ASD heterogeneity (for example, ventriculomegaly with versus without ASD (Kyriakopoulou et al., 2023)), our focus was on fetuses without overt structural brain abnormalities to investigate whether ASD-related differences in brain morphology may nevertheless be detectable by quantitative analysis. Using motion-corrected fetal MRI reconstruction and surface-based morphometry, we quantified whole-brain and regional inner cortical surface area (or *white matter* surface area) during the second and third trimesters and compared these measures to typically developing controls. We tested whether ASD is associated with alterations in cortical surface area, whether such effects show regional heterogeneity across lobes, and whether findings remain robust to variability in image quality and acquisition characteristics inherent to retrospective fetal imaging. Together, this study evaluates whether quantitative fetal MRI-based surface morphometry might detect prenatal cortical differences associated with later ASD.

## 2. Materials and methods

### 2.1. Cohort characteristics

This study uses a retrospective cohort of fetal MRI data acquired at Boston Children’s Hospital between 2014-2024 under IRB approval (P00040121, P00008836) (SI-Figure 1). For the ASD group, approximately 1,283 subjects were initially identified through systematic electronic medical record screening based on ICD-9-CM (299), ICD-10-CM (F84.0) diagnostic codes. Within this cohort, 52 ASD cases with available fetal MRI data were identified and retrieved for further review. Following automated reconstruction and quality assessment procedures, 50 subjects were confirmed to have usable fetal MRI data. To reduce heterogeneity associated with clinically referred imaging, the present study focused on 15 ASD fetuses with no prenatal and postnatal evidence of major structural brain abnormalities from radiologists’ qualitative visual inspection, scanned at a mean gestational age (GA) of 26.7 weeks (range: 20.4-36.3 weeks; 11 (77.3%) male). Of note, although all included ASD fetuses had clinically interpreted normal brain MRI examinations, fetal MRI had been performed following prenatal ultrasound with referral indications including: central nervous system findings (e.g., inferior cerebellar abnormality, prominent cavum vergae, suspected interhemispheric cyst, possible Dandy–Walker variant, or non-visualization of the cavum septum pellucidum), as well as extracranial abnormalities involving the cardiovascular, thoracic, gastrointestinal, renal, craniofacial, and other organ systems (SI-Figure 1C). The remaining ASD cases (N=35) exhibited heterogeneous structural brain malformations and were excluded from the present analyses (SI-Figure 1C).

For cohort characterization, demographic, maternal, perinatal, and environmental variables previously associated with ASD risk (e.g., sex, socioeconomic indicators, maternal medical conditions, substance exposure and birth complications) were extracted from medical records when available (Edlund et al., 2026). Maternal records indicated incidence of anxiety (N=5), autoimmune disorders (N=2), diabetes (N=1), hypertension (N=1), and exposure to drugs (N=2), alcohol (N=2), and smoking (N=1). Neurodevelopmental and psychiatric comorbidities within the ASD group known at the time of manuscript preparation are detailed in SI-Figure 1C. Unfortunately, we did not have access to specific ASD symptom severity measures and related neurodevelopmental assessments such as Autism Diagnostic Observation Schedule, Second Edition (ADOS-2) assessments (Lord et al., 2012) for most of the included subjects, but 3 were flagged as severe, 5 as moderate, and 1 as mild-to-moderate forms of ASD in their medical records.

A control group of 60 typically developing singleton fetuses (TD) was selected from previously presented research fetal imaging cohorts (Gondová et al., 2026; Rollins et al., 2021). Inclusion criteria for the TD group required absence of known brain, body, or genetic abnormalities, and absence of serious maternal medical conditions. TD fetuses were scanned at a mean GA of 28.4 weeks (range: 22.0-35.7 weeks; 34 (56.7%) male). Typical neurodevelopment was confirmed in a subset of TD participants (TD_Optimal_; N = 33) at approximately 2 years of age using the Bayley Scales of Infant and Toddler Development (Bayley, 2006), defined as composite cognitive, language, and motor scores >70. Select demographic and perinatal characteristics of the ASD and TD cohorts are summarized in Table 1.

**Table 1.** Select subject characteristics. Differences between ASD and TD groups for continuous variables were assessed using t-tests with Welch’s correction for unequal variances (t: t-statistic, dof: degrees of freedom, p: p-value, D: Cohen’s D). For categorical variables, differences were evaluated using Chi-squared (Χ²) tests (Χ²: Χ²-statistic, V: Cramer’s V). N: number of subjects.

|  | ASD (N=15) | TD (N=60) |  |
| --- | --- | --- | --- |
| GA at scan (weeks): mean [range] | 26.69 [20.4,36.3] | 28.44 [22.0,35.7] | t=-1.17, dof=15.94,<br>p=0.258, D=0.49 |
| Males/Females: count (%) | 11 (73.3) / 4 (26.7) | 34 (56.7) / 26 (43.3) | $X^2=0.78$, dof=1,<br>p=0.377, V=0.10 |
| Race/Ethnicity: count (%) | 10 (67) | 60 (100) |  |
| *White* | 6 (60) | 49 (82) |  |
| *Black / African American* | 2 (20) | 3 (5) | $X^2=4.88$, dof=3,<br>p=0.18, V=0.26 |
| *Asian* | 0 (0) | 3 (5) |  |
| *Hispanic* | 2 (20) | 5 (8) |  |
| GA at birth (weeks): mean [range] | 37.93 [34.1,40.2] | 39.21 [34.6,41.7] | t=-2.53, dof=18.32,<br>p=0.021, D=0.86 |
| Birth weight (g): mean [range] | 3401.1 [2600,5000] | 3547.3 [2297,4560] | t=-0.77, dof=18.18,<br>p=0.450, D=0.26 |
| Birth weight (Z*): mean [range] | 0.646 [-1.14,5.18] | 0.412 [-2.37,3.05] | t=0.49, dof=17.02,<br>p=0.632, D=0.18 |
| Mode of delivery: count (%) | 13 (86.7) | 52 (86.7) |  |
| *Vaginal* | 5 (38.5) | 38 (73.1) | X<sup>2</sup>=4.13, dof=1,<br>p=0.042, V=0.25 |
| *C-section* | 8 (61.5) | 14 (26.9) |  |
| Maternal age (years): mean [range] | 30.9 [23,41] | 32.4 [22,40] | t=-1.15, dof=20.76,<br>p=0.264, D=0.34 |

\*birth weight z-scores were derived from the normative tables reported by Nicolaides et al., 2018.

### 2.2. MRI acquisition and reconstruction

ASD fetal MRI data was acquired as part of clinical care, resulting in heterogeneity in acquisition protocols. 12 scans were obtained with 3T Siemens systems using multi-planar T2-weighted HASTE sequences optimized for fetal imaging, with parameters included TR=1300-1600ms, TE=70-90 ms, 1 mm in-plane resolution, and ≈4 mm 83slice thickness. 3 scans were acquired on 1.5T GE systems using single-shot fast spin echo (SSFSE) sequences with TR=1300-1600ms, TE=70-90 ms, 1 mm in-plane resolution, and ≈4 mm slice thickness. TD group was derived from our previously reported works (Rollins et al., 2021) and all scans were acquired using a standardized research protocol on either a 3T Siemens MAGNETOM Skyra or Prisma systems with HASTE sequences (TR/TE=1400-1600/99-132ms, 1mm in-plane resolution, 2-3.5 mm slice thickness, field of view adapted to fetal and maternal size (Rollins et al., 2021).

All fetal MRIs underwent identical processing using a validated pipeline including brain extraction (Hong et al., 2020), N4 bias field correction (Tustison et al., 2010), and slice-to-volume reconstruction with NeSVoR (Xu et al., 2023), resulting in super-resolved motion-corrected 3D T2w volumes at 0.5mm isotropic resolution. Details on quality of available data and resulting reconstructions are provided in SI-Figure 2.

### 2.3. Cortical plate (CP) segmentation and inner CP surface extraction

Segmentation of the CP and remaining supratentorial tissues (the “other” label, including non-CP cortex, ventricles, and deep gray matter) was performed using an automated ensemble of 2D U-Net models trained independently on sagittal, coronal, and axial planes. Predictions from the three views were aggregated into 3D volumes, and final labels were assigned based on combined voxel-wise probabilities across models to improve segmentation robustness (Hong et al., 2020). The resulting segmentations were then assessed and manually refined by trained staff based on underlying T2-weighted images. Inner CP surfaces were then extracted using the CIVET marching cubes algorithm (Lepage et al., 2017) using the ChRIS pl-fetal-surface-extract plugin (Zhang, 2025) and post-processed with Taubin smoothing (Taubin, 1995) to suppress overfitting to the coarse voxel boundaries. Surfaces were resampled to standardized meshes of 81,920 triangles and 40,962 vertices, and their quality evaluated both visually and quantitatively via smoothness error (mean curvature difference between each vertex and its neighbors) and boundary distance error (Euclidean distance of vertex to nearest boundary voxel) (SI-Figure 3).

### 2.4. Parcellation of regions of interest

Regional analyses were performed at the lobar level to increase robustness to variability in fetal image quality between TD and ASD cohorts. Major anatomical boundaries separating cerebral lobes could be identified more reliably than finer sulcal- and gyral-based subdivisions, reducing susceptibility to residual segmentation, registration, and parcellation uncertainty while preserving sensitivity to large-scale spatial differences.

To generate lobar parcellations, we employed a previously proposed fetal-adapted surface-based atlas derived from the original 34-region Desikan-Killiany atlas (Desikan et al., 2006) and adapted for fetal neuroanatomy (Vasung et al., 2020). This atlas consists of 21 bilateral regions manually defined on a 29-week fetal surface template (Yun et al., 2019). For the present study, these parcels were combined into six bilateral lobes (frontal, parietal, occipital, temporal, limbic, and insular; SI-Figure 4A) and projected onto individual inner cortical plate surfaces using spherical surface-to-surface registration (Boucher et al., 2009; Robbins, 2004).

### 2.5. Quantification of brain morphometric measures

All measurements were computed in native space. Whole-brain volume was the sum of CP and ‘other’ supratentorial labels multiplied by voxel volume, CP volume was the number of CP voxels multiplied by voxel volume, and inner CP surface area was the total area of triangles on the bilateral inner CP surface, excluding the cingular pole. Lobar inner CP surface area was summed triangle areas within the lobe.

### 2.6. Statistical analysis

#### 2.6.1. Whole-brain and lobar ASD-TD differences

Whole-brain volume, CP volume, and inner CP matter surface area were analyzed using general linear models. Log-transformed morphometric measures were modelled as a function of group (ASD vs TD), sex, and GA. GA was modelled using natural splines to account for potential non-linear developmental effects. To determine the optimal degrees of freedom, we compared models with increasing spline flexibility (linear through fourth-order) using AIC and BIC as primary model selection criteria, with likelihood ratio tests used for confirmation of selection of the best balance between goodness of fit and parsimony. The interaction between group*sex was included only in the whole-brain volume models as it did not show significant relationships to whole-brain CP volume or inner CP surface area and was thus removed for simplicity. Type II Wald F-tests were used to evaluate main effects and interactions. Effect sizes were reported as partial eta squared (η²). Post hoc pairwise comparisons between groups (and sex, where appropriate) were conducted using model-derived estimated marginal means. For interpretability, model-derived estimates are presented on the back-transformed scale, with 95% confidence intervals calculated from the fixed-effects model estimates. All statistical tests were two-sided with α = 0.05. Multiple comparisons were controlled using the false discovery rate (FDR) procedure; additional details regarding correction procedures are provided alongside the corresponding results.

#### 2.6.2. Lobar surface area analysis

Lobar surface area was analyzed using linear mixed-effects models. (Note, lobar volumes were not analyzed as whole-brain CP volume did not show significant group differences). Log-transformed surface area was modelled as a function of group (ASD vs TD), hemisphere, lobar region, and their interaction, and GA (modelled using 3rd order splines with order determined as described in Section 2.6.1). A random intercept for subject was included to account for within-subject correlation across hemispheres and lobes. Significance of fixed effects and interactions was assessed using Type III Wald F-tests. To characterize regional heterogeneity in ASD-related inner CP surface area differences, we then examined the group-by-lobe interaction derived from these linear mixed-effects model. To further quantify the relative structure of the observed regional effects, we derived pairwise interaction contrasts comparing ASD–TD effect sizes between all lobe pairs. These contrasts form a signed, antisymmetric matrix summarizing the extent to which ASD-related deviations in surface area differ across cortical regions. Positive values indicate that the ASD–TD difference is greater in one lobe relative to another (column vs row), whereas negative values indicate the opposite ordering.

#### 2.6.2. Sensitivity analyses

To assess the robustness of observed group effects, we conducted two levels of sensitivity analyses: (i) primary models were repeated after restricting the control group to the neurodevelopmentally confirmed TD_Optimal_ subset. And (ii), we accounted for differences in image quality across subjects using two complementary approaches. In ii-1, models were extended to include the mean quality of image stacks used for reconstruction and the number of stacks as additional covariates to account for potential differences in acquisition quality between ASD and TD cohorts. In ii-2, an alternative propensity score weighting approach was applied using inverse probability weighting to address imbalance in acquisition-related covariates between groups. Propensity scores were estimated from image quality metrics and stack numbers and incorporated into the regression framework. Attenuation or persistence of group effects across sensitivity analyses was used to interpret the robustness of the morphometric findings. (Note, we opted not to perform the analysis with propensity weighting on the lobar level as we cannot confirm assumption that image quality effects are the same across lobes).

#### 2.6.4. Deviations from the typical development

To quantify regional deviations from typical inner CP surface area development, we also constructed a normative model using only the TD cohort. Similar linear mixed-effects model as in Section 2.6.3 (between log-transformed inner CP surface area and hemisphere and lobes, including their interaction, with random intercept for subjects included to account for repeated measurements) was estimated only in the TD subjects. To account for developmental and acquisition-related variability, GA was again modeled using 3rd order splines, and image quality metrics (mean quality and number of stacks) were included as covariates. This model defined expected inner CP surface area conditional on developmental stage, anatomy, and imaging quality. Normative predictions were then generated for all participants (TD and ASD) using fixed effects only. Regional deviation scores were computed as residuals between observed and model-predicted log surface area. These normative deviation scores were then used as the dependent variable in downstream analyses restricted to the ASD cohort. Specifically, linear mixed-effects models were fitted with hemisphere and lobar region as fixed effects and a random intercept for subject, allowing assessment of whether deviation magnitude varied systematically across cortical regions. Given the limited sample size in the ASD cohort (N=15), inferential analyses of these deviations need to be interpreted cautiously and treated as exploratory, alongside descriptive summaries of their overall distribution. Additionally, rank-based statistics were computed to describe the relative lobar ordering of deviation of magnitude across subjects, providing a complementary view of the spatial structure underlying the group-level contrast patterns analyzed in the previous section.

## 3. Results

### 3.1. Whole-brain differences between ASD and TD fetuses

Both groups showed expected developmental increases with GA (F=1385.16, p<0.001, η²=0.98) and effects of sex (F=16.03, p<0.001, η²=0.19) on whole-brain volumes (Table 2A). Additionally, we observed significant group*sex interaction (F=4.69, p=0.034, η²=0.06). Post-hoc comparisons confirmed expected sex difference in the TD group, with females showing significantly lower volume than males (β=−0.09, p<0.001; −8.6%). ASD males and TD males showed significant differences in whole-brain volumes (β=−0.08, p=0.033; −7.7%) that persisted after FDR correction. No group differences were observed in females (SI-Table 1A). Interestingly, we did not observe whole-brain volume differences between males and females in the ASD group (β=0.01, p=812; +1.0%), suggesting a reduction of typical sexual dimorphism in ASD.

**Table 2A.**
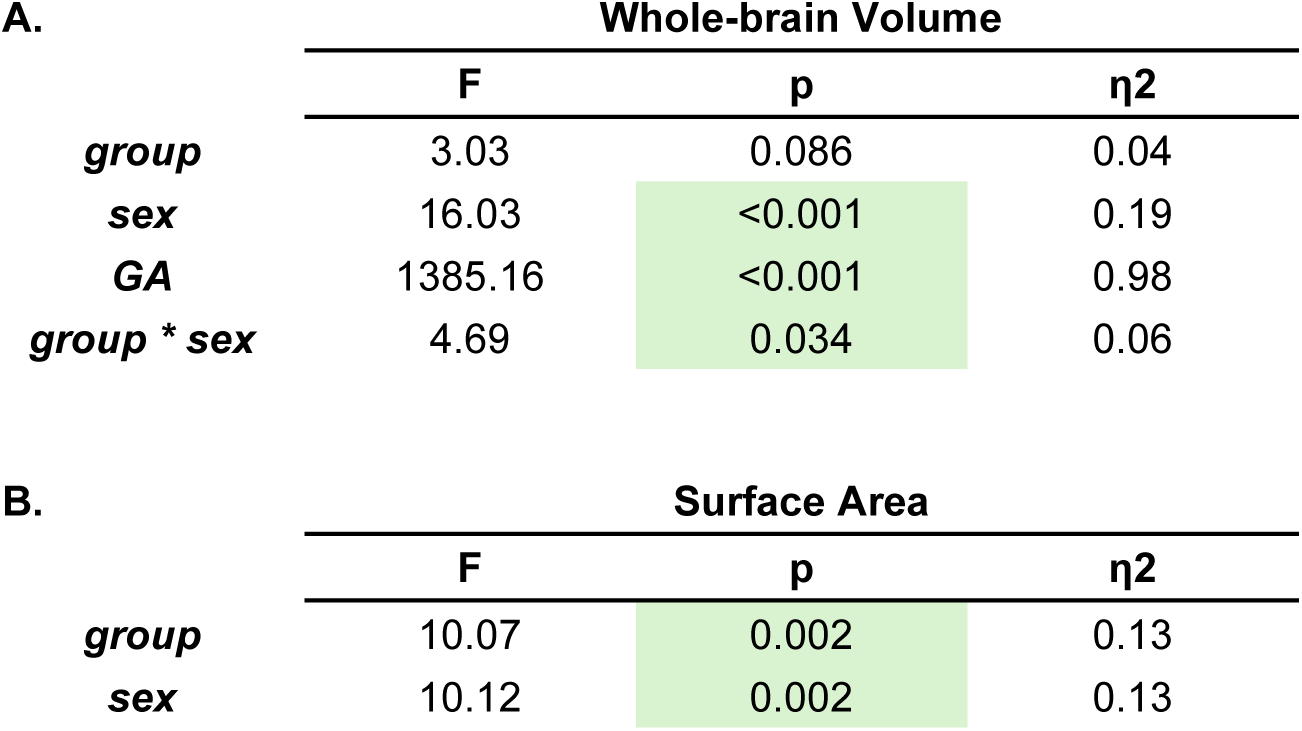
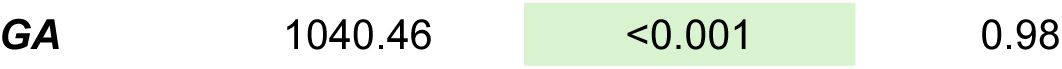
Type II ANOVA results from the linear model of log-transformed whole-brain volume including group, sex, GA (modeled with 2nd order splines), and their interaction. Reported are F-statistics (F), associated p-values (p), and partial eta squared (η²). C. Same as A but for surface area (GA; modeled with 3rd order splines). Note, group*sex was not significant and removed to simplify the model.

We did not observe a similar group difference in whole-brain CP volume (SI-Table 1B). For the whole-brain surface area, we again observed the expected developmental effects of GA (F=1040.46, p<0.001, η²=0.98) and an effect of sex (F=10.12, p=0.002, η²=0.13). Additionally, ASD and TD groups also differed (F=10.07, p=0.002, η²=0.0.13, no significant group*sex interaction) (Table 2B), with ASD-TD contrast (β=-0.08+0.02SE, t ratio=-3.17, p=0.002; −7.7%). In other words, adjusted predicted means (at GA=28.1w) is 8783 mm2 [8608-8961, 0.95% CI] while only 8131 mm2 [7723-8560, 0.95% CI] for ASD subjects.

To assess the impact of potential developmental heterogeneity in the TD group, analyses were repeated in a subset of TD_Optimal_ subjects (SI-Table 2). The results were broadly consistent with the primary analysis. For whole-brain volume, no significant main effect of group (F=2.21, p=0.144, η²=0.05). Group*sex interaction was attenuated (F=2.85, p=0.098, η²=0.06). For surface area, a significant group effect persisted (F=5.57, p=0.023, η²=0.11). These findings indicate that surface area differences might be robust to stricter definitions of the TD group, whereas volume differences remain limited. Note, because CP volume did not show significant group effects in the primary models, sensitivity analyses were not pursued for this measure.

Given significant group differences in acquisition image quality (SI-Table 3, SI-Figure 5), sensitivity analyses were performed: i, including mean HASTE quality and number of stacks as covariates; and ii, using a propensity weighting approach. Including quality metrics as covariates, group effects observed in the primary models were attenuated for both the whole-brain volume (group: F=0.13, p=0.718, η²=0.00, group*sex: F=2.22, p=0.141, η²=0.03), and surface area (group: F=0.44, p=0.508, η²=0.00). QC metrics showed modest associations with the measurements (F=3.09-9.02, p=0.004-0.141, η²=0.03-0.12) (SI-Table 4). The attenuation of observed group effects thus suggests that part of the group-related variance in morphometric measures is not independent of acquisition context and may operate through incidental image quality differences between the groups likely related to clinically *vs* research data acquisition protocols. To overcome this limitation, we further investigate using a modelling approach that included propensity weighting during the model estimation to balance QC distribution between groups. The whole-brain effects of group were still attenuated (group: F=3.41, p=0.069, partial η²=0.05; group*sex: F=1.74, p=0.191, partial η²=0.02), but for surface area, groups effect was significant (F=12.49, p<0.001, partial η²=0.15) (SI-Table 5).

The divergence across whole-brain volume, CP volume, and surface area suggests differential sensitivity of these metrics to biological and acquisition-related factors. Given that surface area showed the most consistent directionality across primary and sensitivity analyses, despite attenuation under some QC-adjusted models, subsequent analyses focus on this measure.

### 3.2. ASD-TD differences of lobar surface area

To evaluate lobar surface area differences between ASD and TD group, we performed mixed-effects modelling. As expected from the whole-brain analyses, group effect was again significant (F=10.11, p=0.001, partial η²=0.11). Importantly, we also observed significant interaction between group*hemi (F=4.57, p=0.033, partial η²=0.01), and group*lobe (F=19.31, p=0.002, η²=0.08), although effects sizes were small. The post-hoc comparisons confirmed significant reductions of surface area in all evaluated lobes (t=-3.73--2.30; p=<0.001-0.014; FDR-corrected) (Table 3, visual summary of predicted relative differences is in Figure 1A).

**Figure 1A.**
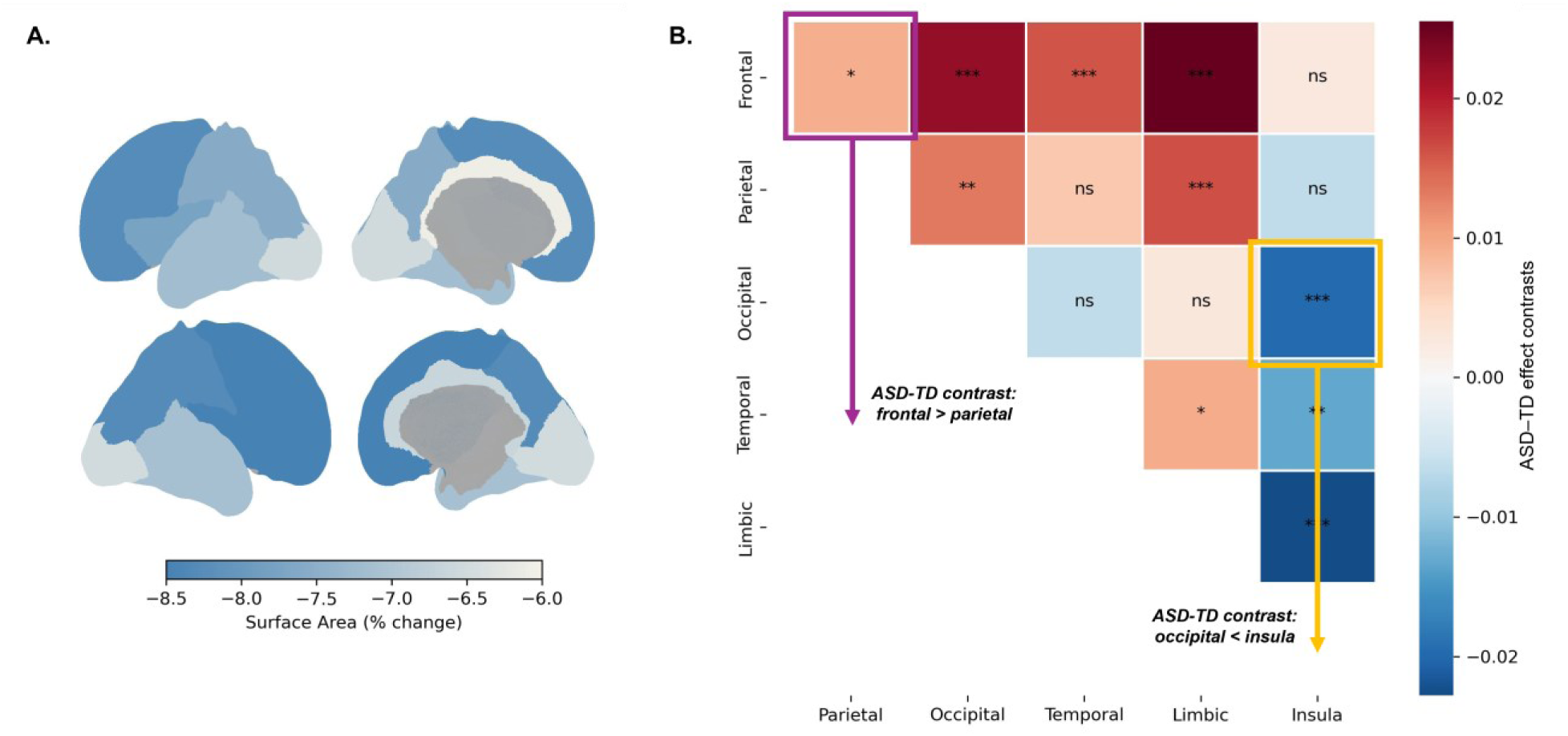
Relative ASD-TD differences summarizing results in Table 3. All Reductions are significant (FDR corrected) **B.** Spatial heterogeneity in group effects from pairwise comparisons. Some lobes have significantly more pronounced surface area reductions in ASD-TD than others (row-column direction, i.e. red in row means given lobe is shows higher reductions than its pair in column, blue means less reduction). Significance codes: p<0.001 (***), p<0.01 (**), p<0.5 (*), p>0.5 (ns); FDR corrected.

**Figure 2A.**
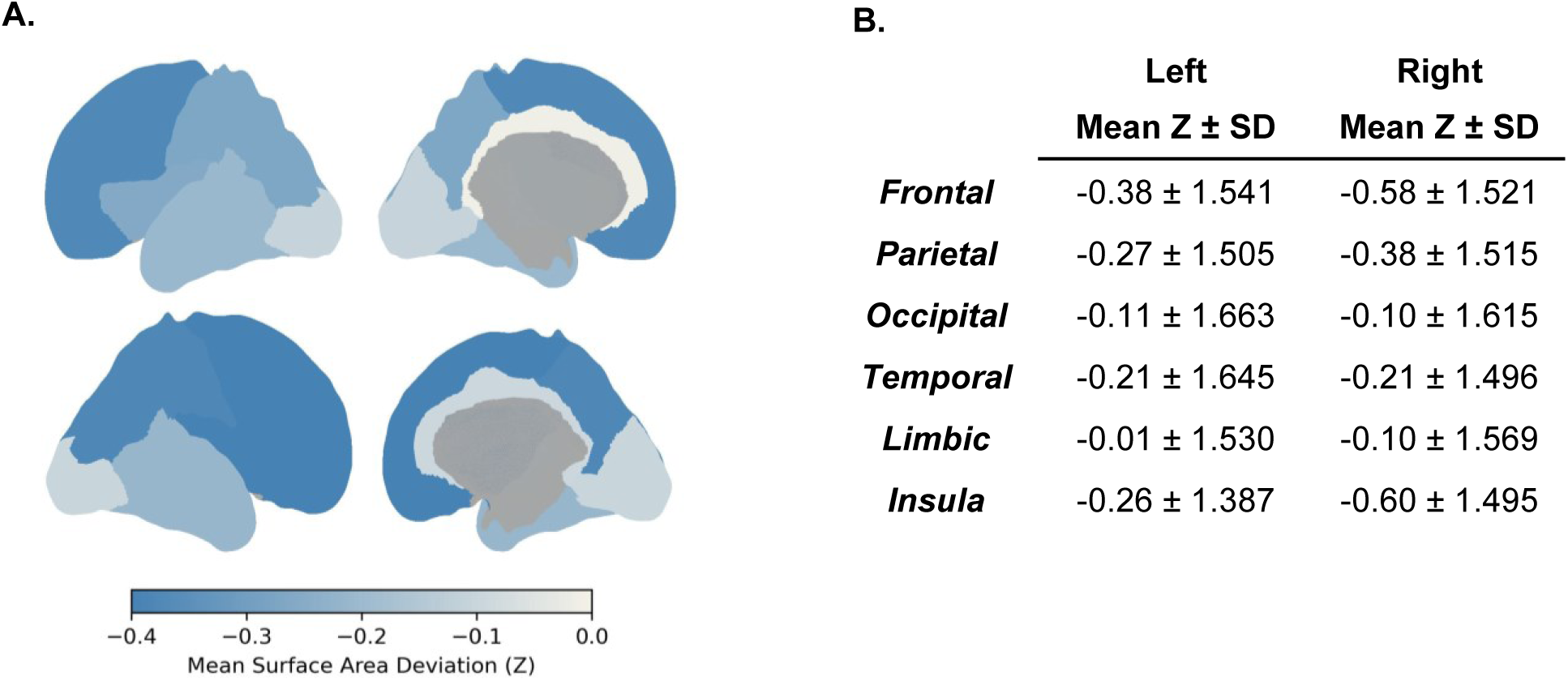
Mean lobar deviations of lobar surface area in ASD group estimated from normative (TD) mixed-effect models (z-scores relative to TD normative model). Normative z-scores were derived from residuals of a TD-trained linear mixed-effects model including lobe, hemisphere, sex, gestational age (3rd-order splines), and imaging quality metrics, with subject-level random intercepts. Note, no deviations were significantly different from 0, but the overall patterns are consistent with group-level magnitudes of the lobar surface area ASD-TD reductions. **B.** Numeric results summarized as ASD mean ± SD per region.

**Table 3.** ASD-TD lobar surface area differences derived from linear mixed-effect modelling of log-transformed surface area, including group, hemisphere, lobe, and their interactions as fixed effects, sex and GA (modeled with 3rd order splines) as covariates, and a random intercept for subject. Group differences were assessed using pairwise contrasts. Reported measures include: estimated marginal means (EMMs; predicted at mean GA=28.1, male sex) and associated 95% confidence intervals (CIs) in the original (back-transformed) scale for interpretability, Wald t-values (t) from log-scale contrasts, associated p-values (p; adjusted for multiple comparisons using FDR method), and effect sizes (d; computed as standardized log-scale mean difference using the standard deviation of log-transformed surface area).

|  |  | ASD [95% CI] | TD [95% CI] | t | p | d |
| --- | --- | --- | --- | --- | --- | --- |
| **Left** | **Frontal** | 14.02 [13.30-14.78] | 15.18 [14.87-15.50] | -3.18 | 0.002 | -0.09 |
|  | **Parietal** | 10.18 [9.66-10.73] | 10.95 [10.73-11.18] | -2.93 | 0.005 | -0.08 |
|  | **Occipital** | 4.60 [4.37-4.85] | 4.90 [4.80-5.01] | -2.53 | 0.014 | -0.07 |
|  | **Temporal** | 7.81 [7.41-8.23] | 8.37 [8.20-8.54] | -2.77 | 0.007 | -0.08 |
|  | **Limbic** | 1.97 [1.87-2.08] | 2.09 [2.04-2.13] | -2.30 | 0.024 | -0.06 |
|  | **Insula** | 2.06 [1.96-2.18] | 2.22 [2.17-2.26] | -2.89 | 0.005 | -0.08 |
| **Right** | **Frontal** | 14.07 [13.35-14.84] | 15.42 [15.11-15.74] | -3.66 | 0.000 | -0.10 |
|  | **Parietal** | 10.13 [9.61-10.68] | 10.97 [10.75-11.20] | -3.18 | 0.002 | -0.09 |
|  | **Occipital** | 4.61 [4.37-4.86] | 4.91 [4.81-5.01] | -2.51 | 0.014 | -0.07 |
|  | **Temporal** | 7.87 [7.46-8.29] | 8.43 [8.26-8.61] | -2.78 | 0.007 | -0.08 |
|  | **Limbic** | 1.96 [1.86-2.07] | 2.09 [2.04-2.13] | -2.50 | 0.015 | -0.07 |
|  | **Insula** | 2.08 [1.97-2.19] | 2.28 [2.23-2.33] | -3.73 | <0.001 | -0.10 |

Pairwise lobar comparisons indicate non-uniform spatial distribution of ASD-related surface area differences across lobes, with modest regional variation in effect magnitude that was not fully stable across sensitivity analyses (Figure 1B).

As before, we performed sensitivity analyses for neurodevelopmentally confirmed TD_Optimal_ subset (SI-Table 7), and mean HASTE quality and number of stacks as covariates. While significant effect of group*hemi (F=18.35, p=0.003, partial η²=0.007) and group*lobe (F=15.51, p=0.008, partial η²=0.08) suggesting spatial differences in ASD effect, the group contrasts across individual lobes were not significant in post-hoc comparisons (SI-Table 8).

### 3.3. ASD-TD differences of lobar surface area (normative)

We further examined regional deviations in inner CP surface area using normative modelling framework trained exclusively on the TD cohort. This model provided expected surface area values conditional on GA, hemisphere, lobar region, sex, and imaging quality metrics, enabling estimation of subject- and lobar-specific deviations relative to typical developmental trajectories. When applied to the ASD cohort, deviation scores showed a small overall negative shift across regions (mean z=-0.27, t=-2.38, p = 0.018), indicating modest reductions in surface area relative to TD-based expectations after accounting for developmental stage and acquisition-related variability. However, no individual lobe-by-hemisphere comparisons survived correction for multiple comparisons, and effect sizes were small with substantial inter-individual variability.

Although regional estimates were consistently shifted in the negative direction, the spatial distribution of deviations did not show statistically robust lobar specificity (SI-Figure 7). Apparent ordering across lobes (e.g., relatively larger deviations in frontal regions compared to posterior regions) should therefore be interpreted cautiously as a complement to group modelling presented in previous sections.

To provide a descriptive summary of spatial structure, we additionally examined within-subject rank ordering of lobar deviations. This confirmed a tendency for lower-ranked deviations in frontal lobe relative to posterior cortices, (grouped across hemisphere, mean ranks: frontal = 3.5±2.36, insula = 4.3±3.04, parietal = 5.3±2.54, temporal = 7.4±2.57, occipital = 9.2±2.51, limbic = 9.3±2.66). However, the proportion of negative deviations was similar across lobes (40–54%), suggesting that the observed spatial pattern reflects *relative differences* in deviation of magnitude across lobes rather than a uniform shift in one direction.

## 4. Discussion

Our study examined prenatal cortical development in fetuses later diagnosed with ASD using fetal MRI and surface-based morphometry. Across multiple modeling strategies, we observed reduced inner cortical surface area in the ASD group relative to typically developing controls during the second and third trimesters. This effect remained evident after restricting analyses to neurodevelopmentally confirmed controls and was more consistent than corresponding measures of whole-brain or cortical plate volume, which showed weaker and less reproducible group differences.

Although the magnitude of surface area differences was modest and partially sensitive to adjustment for acquisition-related variables, including input stack quality and number, the direction of findings remained consistent across sensitivity analyses incorporating imaging quality as covariates and using propensity-based weighting approaches. This suggests that while image quality contributed meaningfully to measurement variability, the observed differences might potentially not be fully explained by systematic differences between clinically acquired ASD scans and research-based TD datasets. Future studies will benefit substantially from prospectively harmonized acquisition protocols and standardized quality-control procedures across groups to corroborate and extend these findings.

At the lobar level, ASD-related differences in surface area were broadly distributed across the cortex, with only modest spatial heterogeneity. Group-by-region effects indicated relatively greater reductions in frontal and insular regions compared to posterior cortices.

Nevertheless, given the modest sample size inherent to this rare fetal ASD imaging cohorts, these regional findings should be interpreted cautiously, particularly for interaction effects where statistical power is limited, and be considered as weak regional modulations of a more distributed effect rather than highly focal lobar abnormalities. Accordingly, the present findings are best interpreted as reflecting subtle regional modulation of a broadly distributed developmental effect rather than highly focal lobar abnormalities.

### 4.1. Cortical surface area and fetal development

The second half of human gestation is characterized by rapid increases in cortical surface area accompanied by progressive cortical folding and increasing regional differentiation of the developing cortex (Bethlehem et al., 2022; Rakic, 1988). During this period, surface area growth becomes a major contributor to overall cerebral expansion and continues to accelerate through the perinatal and early postnatal stages (Garcia et al., 2018; Lyall et al., 2015).

Classical developmental models, including the radial unit hypothesis proposed by (Rakic, 1988), posit that cortical surface area expansion depends predominantly on the number and spatial organization of ontogenetic radial columns generated during corticogenesis, thereby providing a conceptual framework linking macroscopic cortical morphology to the cytoarchitectural organization of the developing brain. Within this framework, tangential cortical expansion emerges from coordinated radial organizational processes, while subsequent work has further emphasized that cortical expansion and its regional heterogeneity reflect not only absolute cell number, but also changes in neuronal morphology, cellular spacing, connectivity, and large-scale cortical architecture (Geschwind & Rakic, 2013).

Interestingly, numerous postmortem studies in ASD have described alterations related to cortical organizational architecture, including altered neuronal morphology, focal disruptions of cortical lamination, and differences in minicolumn organization (Fetit et al., 2021; Hutsler & Casanova, 2016). Although interpretation of these findings continues to be debated, particularly given variability across cortical regions and developmental stages (Buxhoeveden et al., 2006; McKavanagh et al., 2015), many of these observations have been discussed in the context of altered cortical spacing and tangential organizational structure. In particular, (McKavanagh et al., 2015) reported developmental differences in minicolumn width in ASD that appeared more pronounced at younger ages and became less distinct later in development. The authors discussed these findings in relation to prior reports of increased cortical surface area during early childhood ASD that subsequently showed partial convergence toward typically developing trajectories with age (Mak-Fan et al., 2012). Other neuroimaging studies have similarly reported alterations that appear more prominent for cortical surface area than cortical thickness in ASD (Mensen et al., 2017; Ohta et al., 2016), possibly making surface area a particularly informative index of tangential cortical expansion in the context of early ASD-related development.

Importantly, these studies were conducted postnatally, whereas the present study examines surface area differences at an earlier and previously uncharacterized fetal developmental period. Although our observation of reduced cortical surface area in ASD differs in direction from reports of early postnatal surface area expansion, this discrepancy may reflect developmental stage-specific differences in cortical growth trajectories, suggesting that atypical cortical organizational processes may already be present prenatally and continue to evolve across development. Nevertheless, longitudinal information will be required to investigate this question in more depth.

Within this context, the present findings are interesting because they extend investigation of cortical surface area into the prenatal period and suggest subtle reductions of cortical surface area during fetal life. Future studies integrating fetal morphometric characterization with more direct assessments of cortical cytoarchitecture, for example using diffusion imaging, together with genetic risk characterization will allow to further elucidate the neurobiology underlying the developmental trajectories associated with ASD.

### 4.2. Spatial heterogeneity in ASD effects across lobes

At the lobar level, ASD-related reductions in cortical surface area were broadly distributed, with only modest spatial heterogeneity. Nevertheless, the group-by-lobe interactions suggested relatively greater reductions in the frontal lobe and insula compared with posterior cortices, which may be notable in the context of prior developmental models of ASD.

For example, our findings of reduced insular surface area extend prior fetal MRI work reporting increased insular volume in ASD (Ortug et al., 2024), and align with postnatal structural imaging studies implicating the insula across childhood and adulthood (Cauda et al., 2011; Kosaka et al., 2010; Parellada et al., 2017; Parikshak et al., 2013; S. S.-H. Wang et al., 2014; Willsey et al., 2013). Differences in the direction of effects across studies likely reflect sensitivity to distinct morphometric dimensions rather than divergent findings. Cortical volume integrates both surface area and thickness, which are partially dissociable developmental features that may vary independently across development. In this context, increased insular volume alongside reduced surface area likely reflects different aspects of underlying developmental processes, providing a complementary view of insular cortical organization in ASD.

Additionally, transcriptomic studies have implicated early disruption of gene expression programs associated primarily with synaptic development and neuronal differentiation, with enrichment in association cortices including frontal cortex during mid-gestation (Parikshak et al., 2013; S. S.-H. Wang et al., 2014; Willsey et al., 2013). In addition, postnatal neuroimaging studies frequently report involvement of frontal and higher-order associative regions, across multiple modalities including cortical morphology, structural and functional connectivity, and growth trajectories (Courchesne et al.; Ecker et al.). However, these findings are often variable in spatial specificity and directionality, reflecting a broader pattern of heterogeneity across ASD neuroimaging literature (Halliday et al., 2024).

This variability is likely multifactorial, reflecting both biological heterogeneity in ASD and differences in developmental timing across studies. Association cortices, particularly prefrontal regions, follow prolonged maturational trajectories extending from late gestation through early postnatal life (Kostović, 2020), potentially increasing sensitivity to variation in developmental timing and stage-dependent perturbations. In this context, the modest frontal predominance observed here may reflect early divergence in cortical expansion patterns during prenatal development but should be understood within a developmental continuum rather than at isolated developmental time points. Longitudinal approaches spanning prenatal and postnatal (Clairmont et al., 2022; Ecker et al., 2015) stages will be essential to determine how early cortical differences relate to later neurodevelopmental trajectories and outcomes, particularly given the complex interplay of prenatal, perinatal, and postnatal influences contributing to ASD risk, heterogeneity, and clinical presentation (Love et al., 2024; C. Wang et al., 2017). More integrative approaches combining complementary imaging modalities, genetic characterization, and longitudinal developmental trajectories will likely be necessary to better understand the biological mechanisms contributing to ASD at the individual level.

Moreover, fetal MRI reconstruction and surface extraction remain technically challenging, particularly in the presence of fetal motion, variable tissue contrast, and rapidly changing anatomy across gestation. Although all data underwent identical processing pipelines and manual quality assessment, regional analyses were intentionally conducted at the lobar rather than finer gyral level to reduce sensitivity to residual local registration and parcellation inaccuracies. While this necessarily limits anatomical specificity, and consequently more localized cortical alterations cannot be excluded, the relatively small magnitude of the overall lobar differences suggests that the observed ASD–TD differences are better interpreted as reflecting weak spatial modulation of a relatively diffuse effect and conclusions regarding lobar specificity should remain cautious.

## 5. Conclusion

In summary, we report modest reductions in prenatal cortical surface area in fetuses later diagnosed with ASD, detectable in vivo during mid-to-late gestation using fetal MRI. These differences were directionally consistent across multiple analytical frameworks but showed limited regional specificity and partial sensitivity to acquisition-related factors. The present study further demonstrates the feasibility of studying prenatal cortical morphogenesis in fetuses later diagnosed with ASD using advanced fetal MRI reconstruction and surface-based morphometric approaches, providing an initial in vivo characterization of prenatal cortical surface area development associated with later ASD diagnosis.

Taken together, our findings suggest that aspects of cortical morphometric development may diverge subtly prior to birth in individuals later diagnosed with ASD. Given the rarity of available fetal ASD imaging data and the retrospective nature of clinically acquired fetal cohorts, these findings provide a unique in vivo window into early cortical developmental variation associated with later ASD diagnosis. However, the present results should be interpreted as preliminary and primarily descriptive in nature. The retrospective clinically referred cohort introduces potential ascertainment biases associated with indications for fetal imaging, and the available ASD cases likely do not fully represent the broader ASD population. Although subjects with major structural abnormalities were excluded, substantial heterogeneity likely remains with respect to ASD presentation, associated comorbidities, and later neurodevelopmental outcomes. Larger sample sizes, ideally with accompanying genetic data, will be necessary to investigate this heterogeneity in greater detail. Accordingly, these findings underscore both the potential of fetal neuroimaging for investigating early neurodevelopmental variability and the need for replication in prospectively collected, deeply phenotyped, and methodologically harmonized cohorts.

## Acknowledgments

This work was supported by the National Institutes of Health: National Institute of Biomedical Imaging and Bioengineering (R01EB031170) and the National Institute of Neurological Disorders and Stroke (K23NS101120, R01NS114087, and R01NS121334), and the Boston Children’s Hospital (92436). We thank Jason Liu for his invaluable help with patients’ record review. We are deeply grateful to the families whose participation made this research possible.

## Materials and software availability

Image processing code is available on GitHub (https://github.com/FNNDSC). Other processing used publicly available toolboxes detailed in *Materials and Methods*. Analyses were done mainly in Python (3.11.2) and RStudio (4.4.3). Manual corrections and some visualizations used Freeview (3.0).

## Supplementary Information

### 7.1. Cohort characteristics

**SI-Figure 1A.**
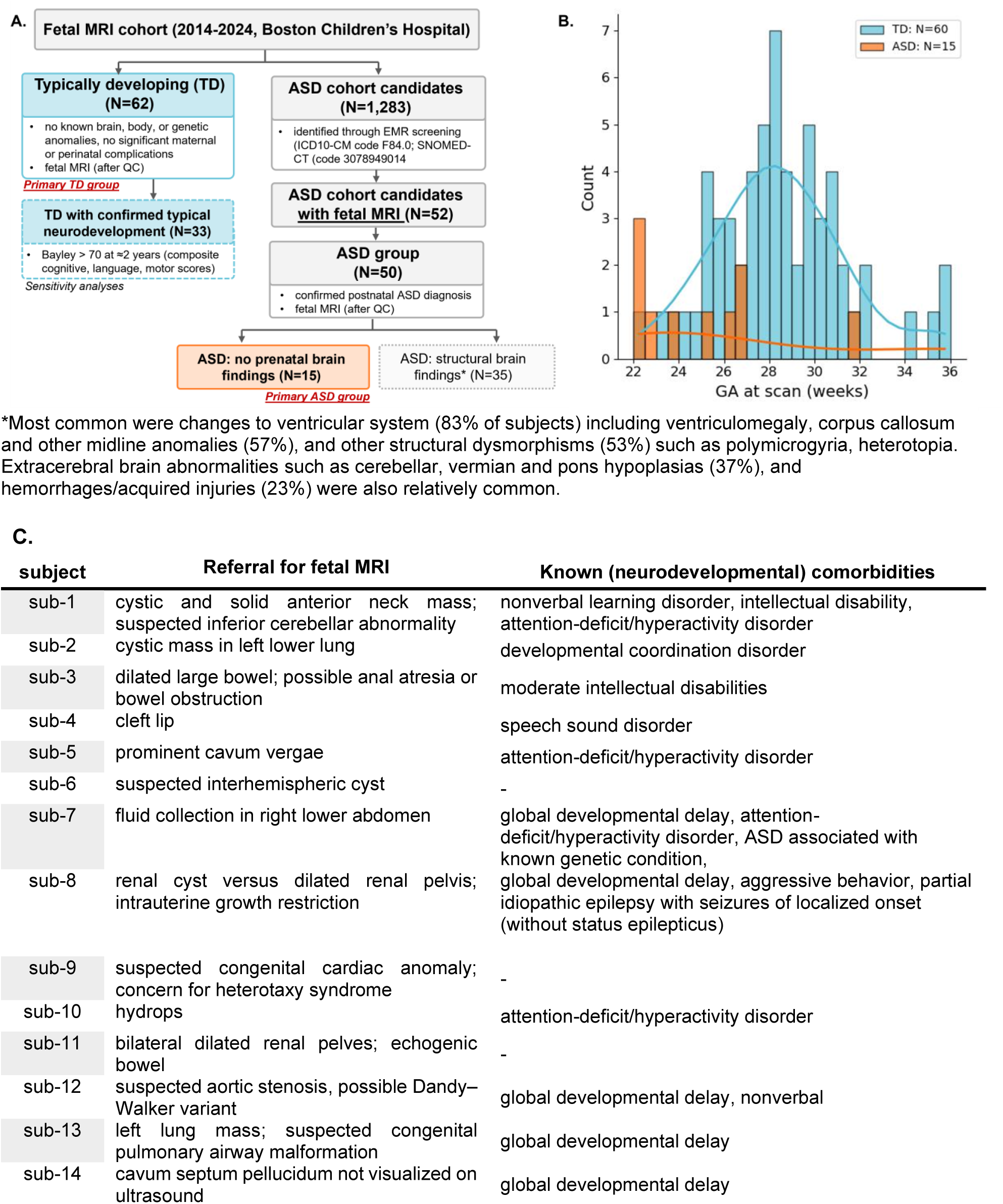
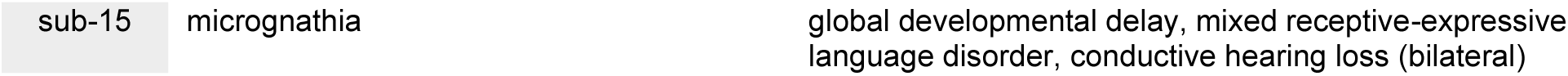
Overview of the cohort creation. **B.** GA distribution of the final subject groups. **C.** Prenatal ultrasound findings prompting fetal MRI referral for each included ASD subject and retrospectively identified neurodevelopmental, psychiatric, and related comorbidities documented in the medical record at the time of analysis (May 2026).

### 7.2. Data quality and processing

**SI-Figure 2A.**
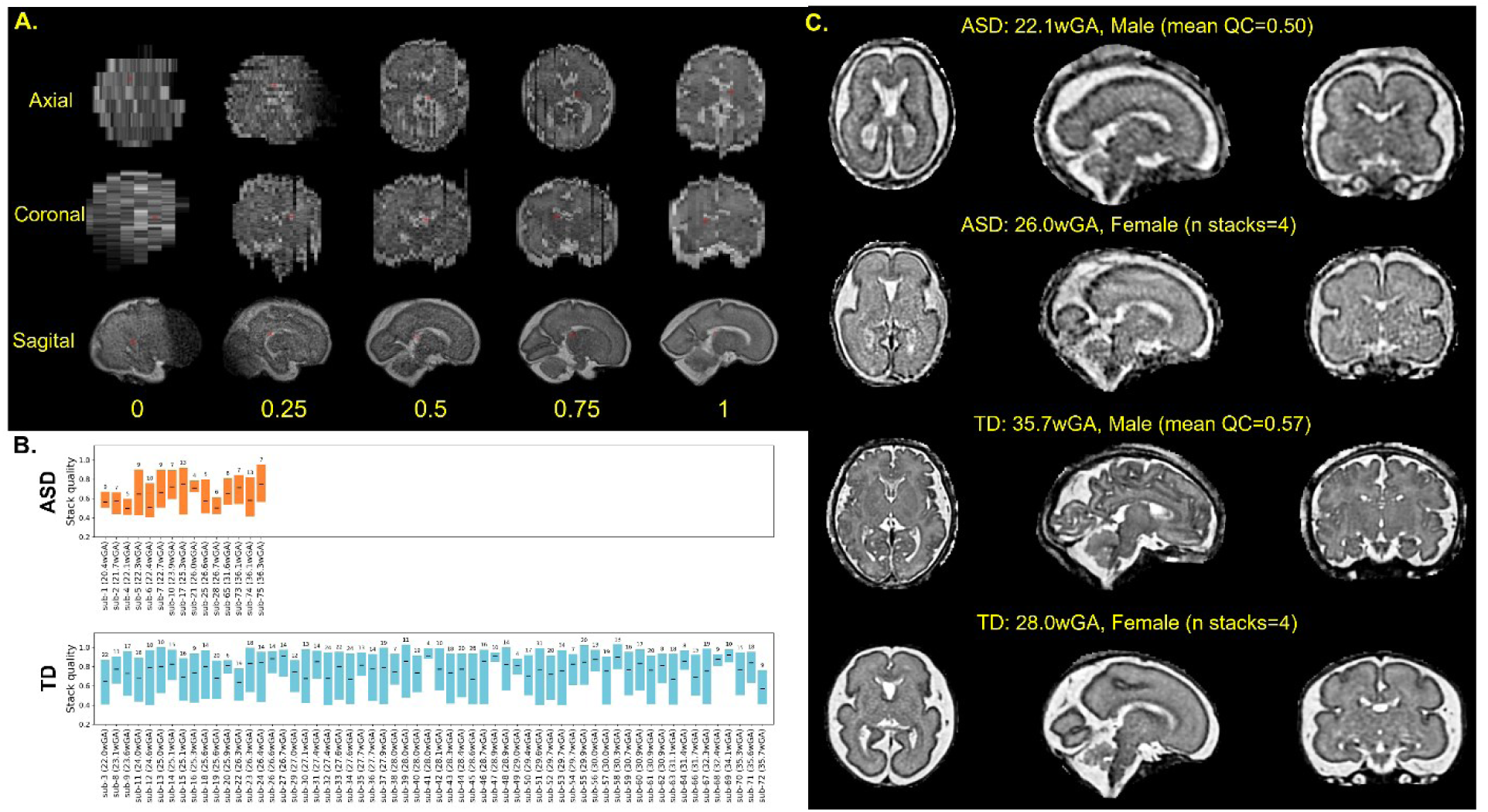
Illustration of the automated stack quality assessment. Before reconstruction, each available image stack was assigned a value between 0-1 (0: very poor, unusable due to severe motion/blurring; 0.25: poor, usable only for basic reconstruction with evident artifacts; 0.5: acceptable, some blurring but preserved anatomical detail; 0.75: good, minimal blurring, clear tissue boundaries; 1: excellent, sharp anatomical detail, negligible motion artifacts) using a deep-learning model trained on slice-level manual quality ratings from two expert readers as ground truth. Only the subjects with at least 3 stacks with quality ≥0.4 were subsequently reconstructed with NeSVoR (this threshold achieves that stacks with excessive motion and generally low quality do not prevent effective reconstruction). **B.** Distribution of stack quality and numbers across included subjects, ordered by GA at scan. Black bars indicate each subject’s mean quality across included stacks; the boxes span the minimum-maximum quality range, and the number above each box shows the final number of stacks used for reconstruction. **C.** Example reconstructions for representative TD and ASD subjects showing subjects with the lowest mean stack quality and subjects with the fewest included stacks.

**SI-Figure 3.**
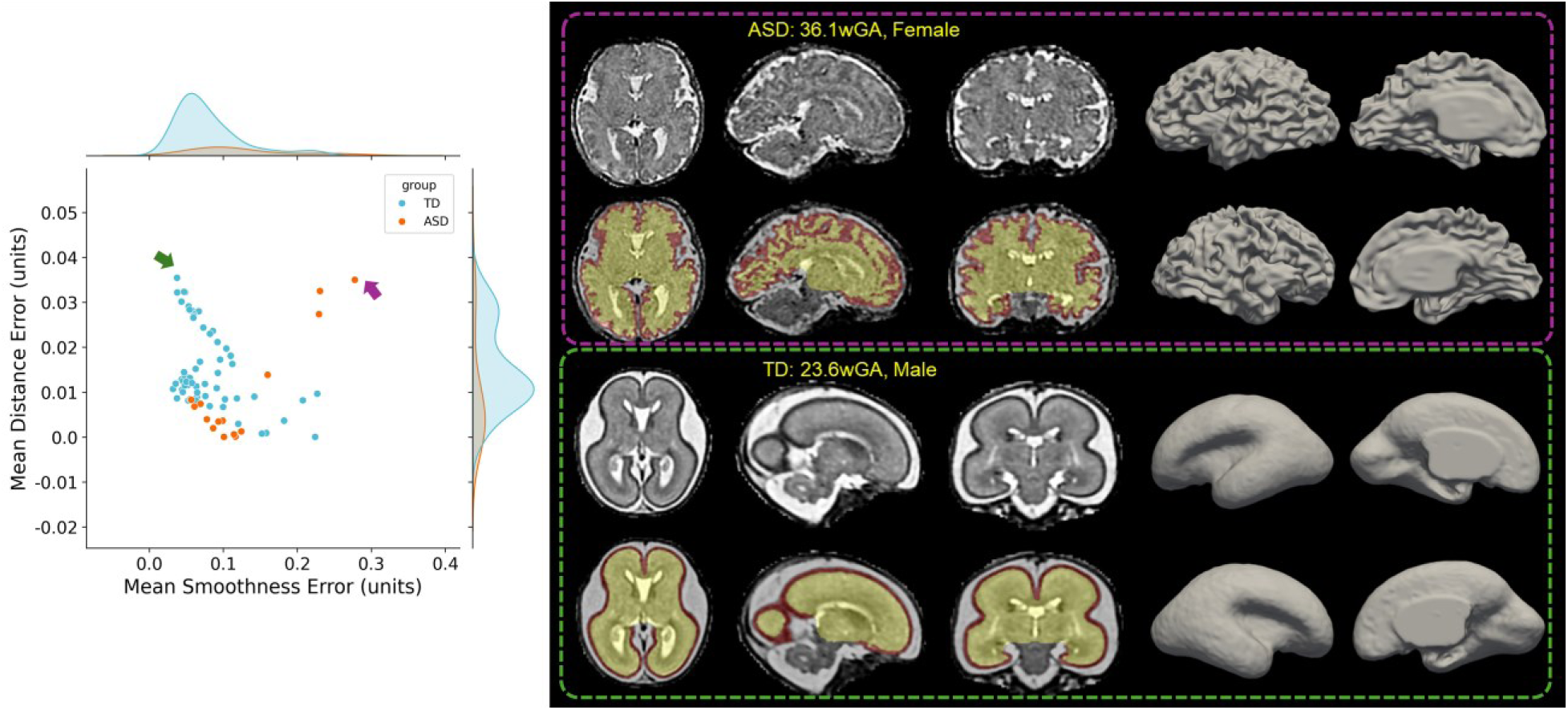
Quality control of surface extraction for the inner CP surface across subjects, together with example T2w reconstruction, their associated tissue segmentation (CP-red, other supratentorial tissue - yellow) in 31w template space for the subjects with the lowest surface extraction quality, as assessed by smoothness (in purple box) and distance errors (in green box).

**SI-Figure 4.**
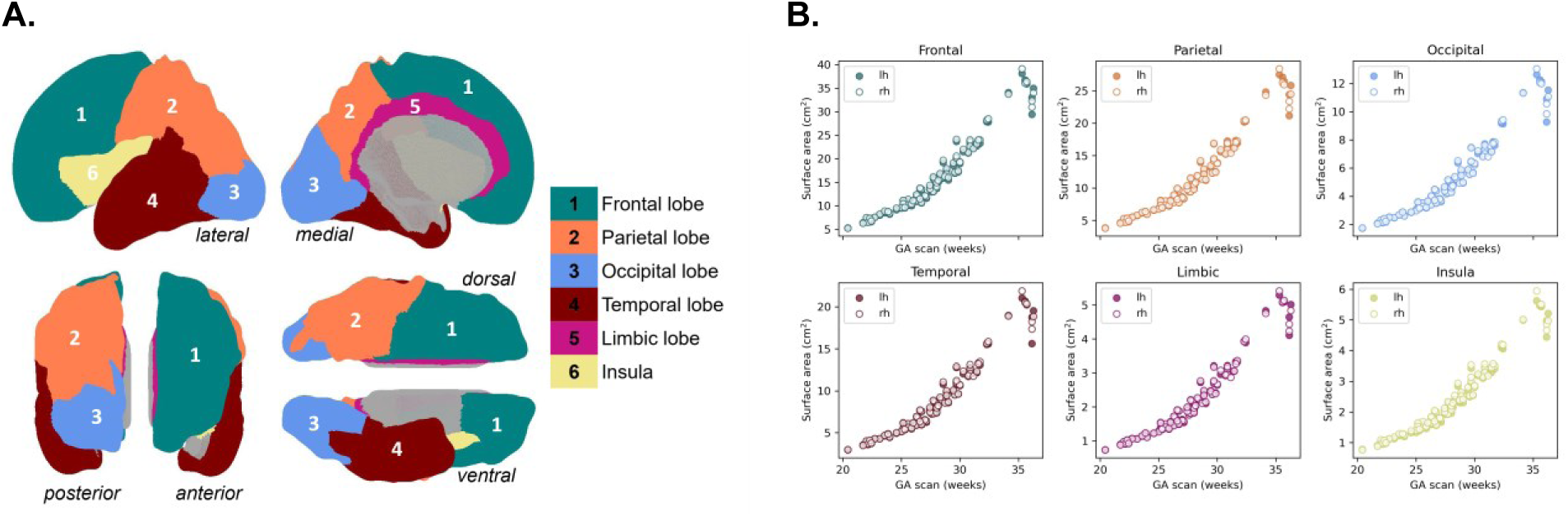
Visualization of the regions for sub-whole-brain level analyses, along with the corresponding color scheme. **B.** Visualization of changes in inner CP surface area with GA showing expected developmental increases.

### 7.3. Whole-brain analyses

**SI-Table 1A.** Post hoc pairwise comparisons of estimated marginal means from the linear model (log-transformed whole-brain volume) across group and sex, adjusted for multiple comparisons using the false discovery rate (FDR). **B.** Type II ANOVA results from the linear model of log-transformed CP volume including group, sex, GA (modeled with 3rd order splines), and their interaction. Reported are F-statistics (F), associated p-values (p), and partial eta squared (η²).

| A. | Estimate ± SE | T (ratio) | p |
| --- | --- | --- | --- |
| *ASD Female – TD Female* | 0.02 ± 0.04 | 0.55 | 0.812 |
| *ASD Female – ASD Male* | 0.01 ± 0.04 | 0.34 | 0.812 |
| *ASD Female – TD Male* | -0.06 ± 0.04 | -1.66 | 0.205 |
| *TD Female – ASD Male* | -0.01 ± 0.03 | -0.24 | 0.812 |
| **TD Female – TD Male** | -0.09 ± 0.02 | -4.53 | <0.001 |
| **ASD Male – TD Male** | -0.08 ± 0.03 | -2.62 | 0.033 |

| B. | CP V |  |  |
| --- | --- | --- | --- |
|  | F | p | η<sup>2</sup> |
| **group** | 0.39 | 0.534 | 0.00 |
| **sex** | 5.02 | 0.028 | 0.07 |
| **GA** | 866.38 | <0.001 | 0.97 |
| **group \* sex** | 0.017 | 0.898 | 0.00 |

#### Sensitivity analyses

##### 1. TD: confirmed typical neurodevelopment subset (N=36)

**SI-Table 2A.** Type II ANOVA results from the linear model of log-transformed whole volume including group (TD restricted to subjects with postnatal neurodevelopmental follow-up), sex, GA (modeled with 2nd order splines), and their interaction. Reported are F-statistics (F), associated p-values (p), and partial eta squared (η²). **B.** Same as A but for surface area (GA modeled with 3rd order splines). Note, as group*sex was not significant in primary models, it was not included here.

| A. | Whole-brain Volume |  |  |
| --- | --- | --- | --- |
|  | F | p | η<sup>2</sup> |
| **group** | 2.21 | 0.144 | 0.05 |
| **sex** | 5.36 | 0.025 | 0.11 |
| **GA** | 973.69 | <0.001 | 0.98 |
| **group \* sex** | 2.85 | 0.098 | 0.06 |

| B. | Surface Area |  |  |
| --- | --- | --- | --- |
|  | F | p | η<sup>2</sup> |
| **group** | 5.57 | 0.023 | 0.11 |
| **sex** | 3.70 | 0.061 | 0.08 |
| **GA** | 699.12 | <0.001 | 0.98 |

##### 2. Effect of data quality

**SI-Table 3.**
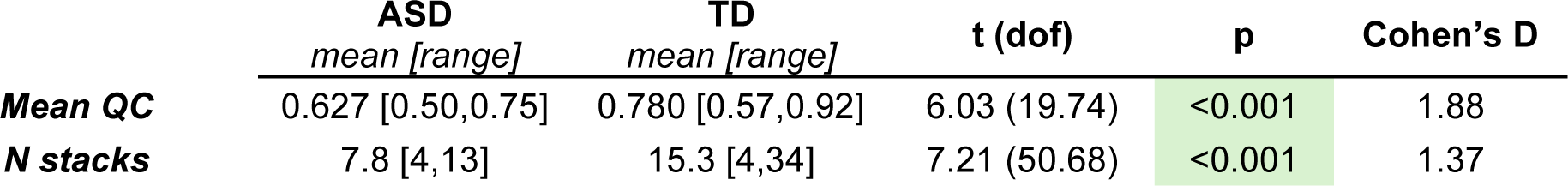
Group differences in image and surface quality between ASD and TD cohorts. Values are reported as mean [range]. Between-group differences were assessed using independent t-tests with Welch’s correction. Reported statistics include t-statistic (t) with corresponding degrees of freedom (dof), p-values (p), and effect sizes (Cohen’s d).

|  | ASD | TD | t (dof) | p | Cohen's D |
| --- | --- | --- | --- | --- | --- |
|  | mean [range] | mean [range] |  |  |  |
| **Mean QC** | 0.627 [0.50,0.75] | 0.780 [0.57,0.92] | 6.03 (19.74) | <0.001 | 1.88 |
| **N stacks** | 7.8 [4,13] | 15.3 [4,34] | 7.21 (50.68) | <0.001 | 1.37 |

###### a. Using image quality as covariates

**SI-Table 4A.** Type II ANOVA results from the linear model of log-transformed whole volume including group, sex, GA (modeled with 2nd order splines), their interaction, and additionally measurements of acquisition quality (mean QC and number of stacks used in reconstruction). Reported are F-statistics (F), associated p-values (p), and partial eta squared (η²). **B.** Same as A but for surface area (GA modeled with 3rd order splines). Note, as group*sex was not significant in primary models, it was not included here.

| A. | Whole-brain Volume |  |  |
| --- | --- | --- | --- |
| | F | p | $\eta^2$ |
| *group* | 0.13 | 0.718 | 0.00 |
| *sex* | 15.36 | <0.001 | 0.19 |
| *GA* | 1346.70 | <0.001 | 0.98 |
| *Mean QC* | 3.54 | 0.064 | 0.05 |
| *N stacks* | 3.09 | 0.083 | 0.04 |
| *group \* sex* | 2.22 | 0.141 | 0.03 |

| B. | Surface Area |  |  |
| --- | --- | --- | --- |
| | F | p | $\eta^2$ |
| *group* | 0.44 | 0.508 | 0.00 |
| *sex* | 12.10 | <0.001 | 0.15 |
| *GA* | 1067.49 | <0.001 | 0.98 |
| *Mean QC* | 9.02 | 0.004 | 0.12 |
| *N stacks* | 3.62 | 0.062 | 0.05 |

###### b. Propensity weighting

In attempt to reduce QC imbalance while preserving the full samples size (as opposed to matching TD quality), we also conducted a sensitivity analysis using propensity score weighting. Propensity scores were estimated from mean QC and number of stacks values using generalized linear models (GLMs, *WeightIt* 1.7.0 R package) and used to derive inverse probability weights (weights were trimmed due to limited overlap between ASD and TD groups because of large data quality differences). Weighted linear models were then re-estimated to assess the robustness of group effects after accounting for differences in image quality.

**SI-Figure 5.**
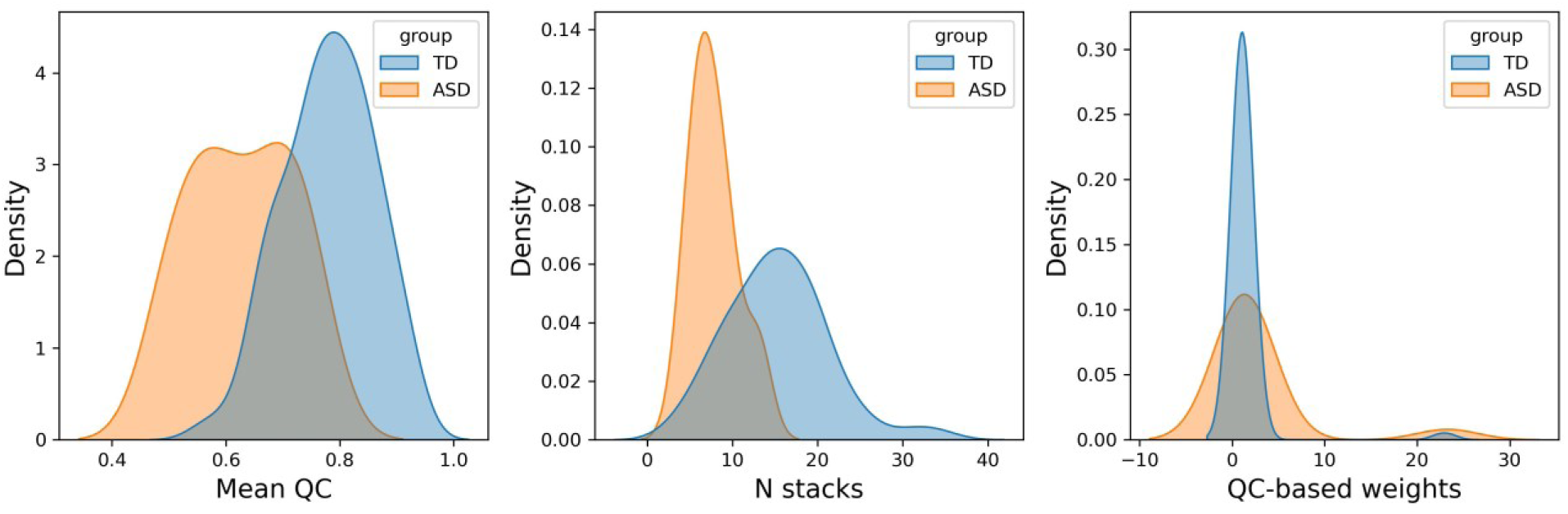
Density plots visualizing group differences in data quality and estimated weights for sensitivity analyses.

**SI-Table 5A.** Type II ANOVA results from the linear model of log-transformed whole volume including group, sex, GA (modeled with 2nd order splines), and their interaction. The observations were weighted based QC metrics. Reported are F-statistics (F), associated p-values (p), and partial eta squared (η²). **B.** Same as A but for surface area (GA modeled with 3rd order splines). Note, as group*sex was not significant in primary models, it was not included here.

| A. | Whole-brain Volume |  |  |
| --- | --- | --- | --- |
| | F | p | $\eta^2$ |
| *group* | 3.41 | 0.069 | 0.05 |
| *sex* | 16.63 | <0.001 | 0.19 |
| *GA* | 1889.18 | <0.001 | 0.98 |
| *group \* sex* | 1.74 | 0.191 | 0.02 |

| B. | Surface Area |  |  |
| --- | --- | --- | --- |
| | F | p | $\eta^2$ |
| *group* | 12.49 | <0.001 | 0.15 |
| *sex* | 15.86 | <0.001 | 0.18 |
| *GA* | 2098.55 | <0.001 | 0.98 |

### 7.4. Lobar analyses

**SI-Table 6.** Type III ANOVA results from mixed-effect models of log-transformed lobar surface area (within-subject error, GA 3rd order spline) Reported are F-statistics (F), associated p-values (p), and partial eta squared (η²).

|  | Surface Area |  |  |
| --- | --- | --- | --- |
| | F | p | $\eta^2$ |
| *group* | 10.11 | 0.001 | 0.11 |
| *hemi* | 0.54 | 0.461 | 0.03 |
| *lobe* | - | - | - |
| *sex* | 10.06 | 0.002 | 0.13 |
| *GA* | 3042.00 | <0.001 | 0.98 |
| *group\*hemi* | 4.57 | 0.033 | 0.01 |
| *group\*lobe* | 19.31 | 0.002 | 0.08 |
| *hemi\*lobe* | 6.16 | 0.291 | 0.05 |
| *group\*hemi\*lobe* | 10.02 | 0.075 | 0.01 |

#### Sensitivity analyses

##### 1. TD: confirmed typical neurodevelopment subset (N=36)

**SI-Table 7.** Type III ANOVA results from mixed-effect models of log-transformed lobar surface area (within-subject error, GA 3rd order spline). Reported are F-statistics (F), associated p-values (p), and partial eta squared (η²).

|  | Surface Area |  |  |
| --- | --- | --- | --- |
| | F | p | $\eta^2$ |
| *group* | 5.72 | <0.001 | 0.10 |
| *hemi* | 0.58 | 0.017 | 0.03 |
| *lobe* | - | - | - |
| *sex* | 3.62 | 0.057 | 0.07 |
| *GA* | 2070.10 | <0.001 | 0.98 |
| *group\*hemi* | 3.12 | 0.077 | 0.01 |
| *group\*lobe* | 15.35 | 0.009 | 0.09 |
| *hemi\*lobe* | 6.56 | 0.255 | 0.06 |
| *group\*hemi\*lobe* | 7.73 | 0.172 | 0.01 |

##### 2. Effect of data quality

###### a. Using image quality as covariates

**SI-Table 8A.**
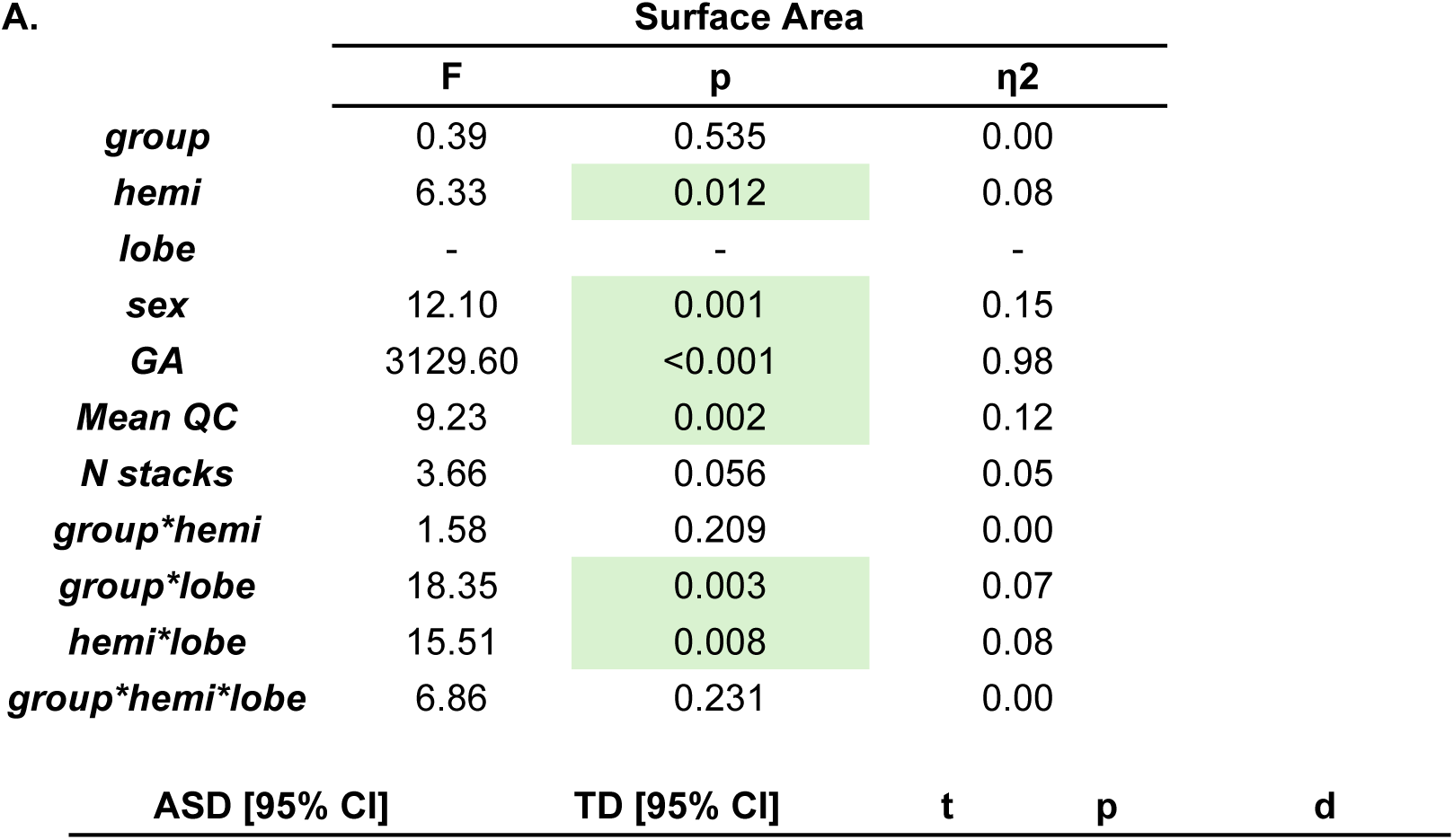
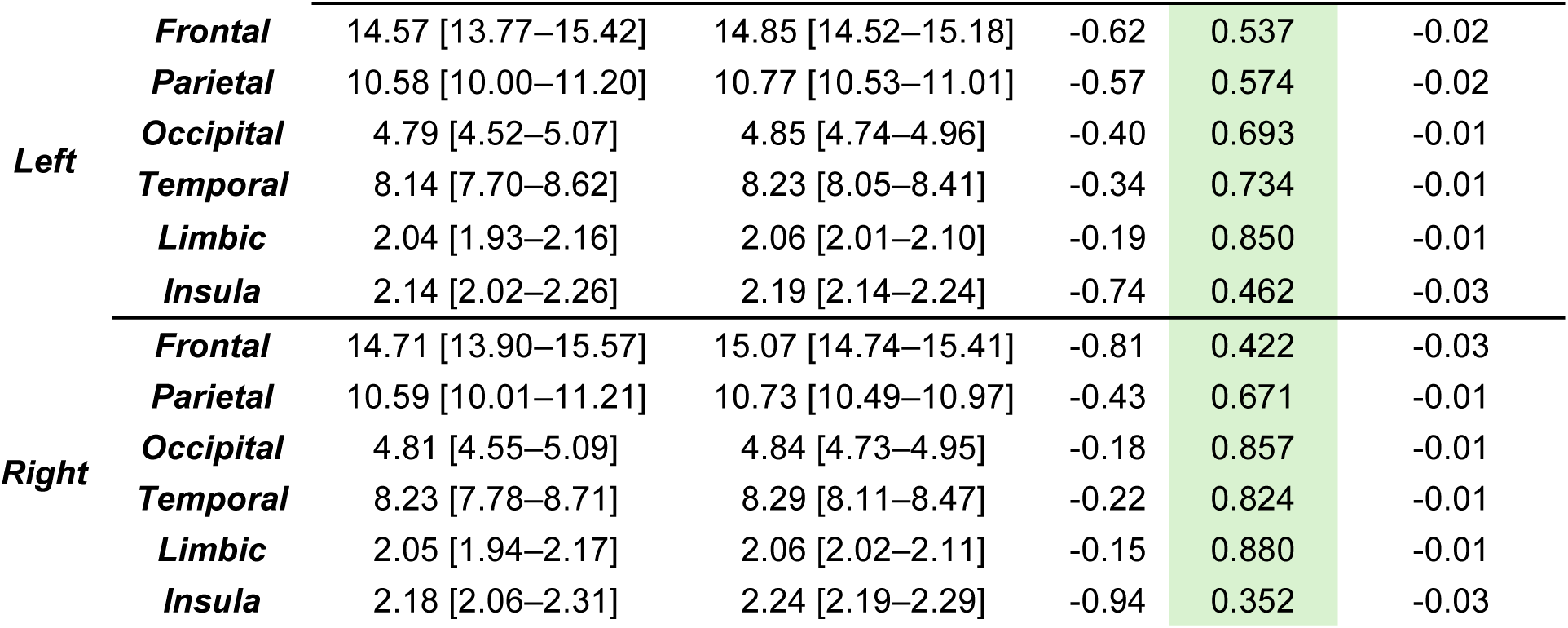
Type III ANOVA results from mixed-effect models of log-transformed lobar surface area (within-subject error, GA 3rd order spline), including measurements of acquisition quality (mean QC and number of stacks used in reconstruction) as covariates. Reported are F-statistics (F), associated p-values (p), and partial eta squared (η²). **B.** ASD-TD lobar surface area differences derived from A. Group differences were assessed using pairwise contrasts. Reported measures include: estimated marginal means (EMMs; predicted at mean GA=28.1, male sex) and associated 95% confidence intervals (CIs) in the original (back-transformed) scale for interpretability, Wald t-values (t) from log-scale contrasts, associated p-values (p; adjusted for multiple comparisons using FDR method), and effect sizes (d; computed as standardized log-scale mean difference using the standard deviation of log-transformed surface area).

| **A.** | Surface Area |  |  |
| --- | --- | --- | --- |
| | F | p | $\eta^2$ |
| *group* | 0.39 | 0.535 | 0.00 |
| *hemi* | 6.33 | 0.012 | 0.08 |
| *lobe* | - | - | - |
| *sex* | 12.10 | 0.001 | 0.15 |
| *GA* | 3129.60 | <0.001 | 0.98 |
| *Mean QC* | 9.23 | 0.002 | 0.12 |
| *N stacks* | 3.66 | 0.056 | 0.05 |
| *group\*hemi* | 1.58 | 0.209 | 0.00 |
| *group\*lobe* | 18.35 | 0.003 | 0.07 |
| *hemi\*lobe* | 15.51 | 0.008 | 0.08 |
| *group\*hemi\*lobe* | 6.86 | 0.231 | 0.00 |

| **B** | ASD [95% CI] | TD [95% CI] | t | p | d |
| --- | --- | --- | --- | --- | --- |

|  |  |  |  |  |  |  |
| --- | --- | --- | --- | --- | --- | --- |
| **Left** | **Frontal** | 14.57 [13.77–15.42] | 14.85 [14.52–15.18] | -0.62 | 0.537 | -0.02 |
|  | **Parietal** | 10.58 [10.00–11.20] | 10.77 [10.53–11.01] | -0.57 | 0.574 | -0.02 |
|  | **Occipital** | 4.79 [4.52–5.07] | 4.85 [4.74–4.96] | -0.40 | 0.693 | -0.01 |
|  | **Temporal** | 8.14 [7.70–8.62] | 8.23 [8.05–8.41] | -0.34 | 0.734 | -0.01 |
|  | **Limbic** | 2.04 [1.93–2.16] | 2.06 [2.01–2.10] | -0.19 | 0.850 | -0.01 |
|  | **Insula** | 2.14 [2.02–2.26] | 2.19 [2.14–2.24] | -0.74 | 0.462 | -0.03 |
| **Right** | **Frontal** | 14.71 [13.90–15.57] | 15.07 [14.74–15.41] | -0.81 | 0.422 | -0.03 |
|  | **Parietal** | 10.59 [10.01–11.21] | 10.73 [10.49–10.97] | -0.43 | 0.671 | -0.01 |
|  | **Occipital** | 4.81 [4.55–5.09] | 4.84 [4.73–4.95] | -0.18 | 0.857 | -0.01 |
|  | **Temporal** | 8.23 [7.78–8.71] | 8.29 [8.11–8.47] | -0.22 | 0.824 | -0.01 |
|  | **Limbic** | 2.05 [1.94–2.17] | 2.06 [2.02–2.11] | -0.15 | 0.880 | -0.01 |
|  | **Insula** | 2.18 [2.06–2.31] | 2.24 [2.19–2.29] | -0.94 | 0.352 | -0.03 |

**SI-Figure 6A.**
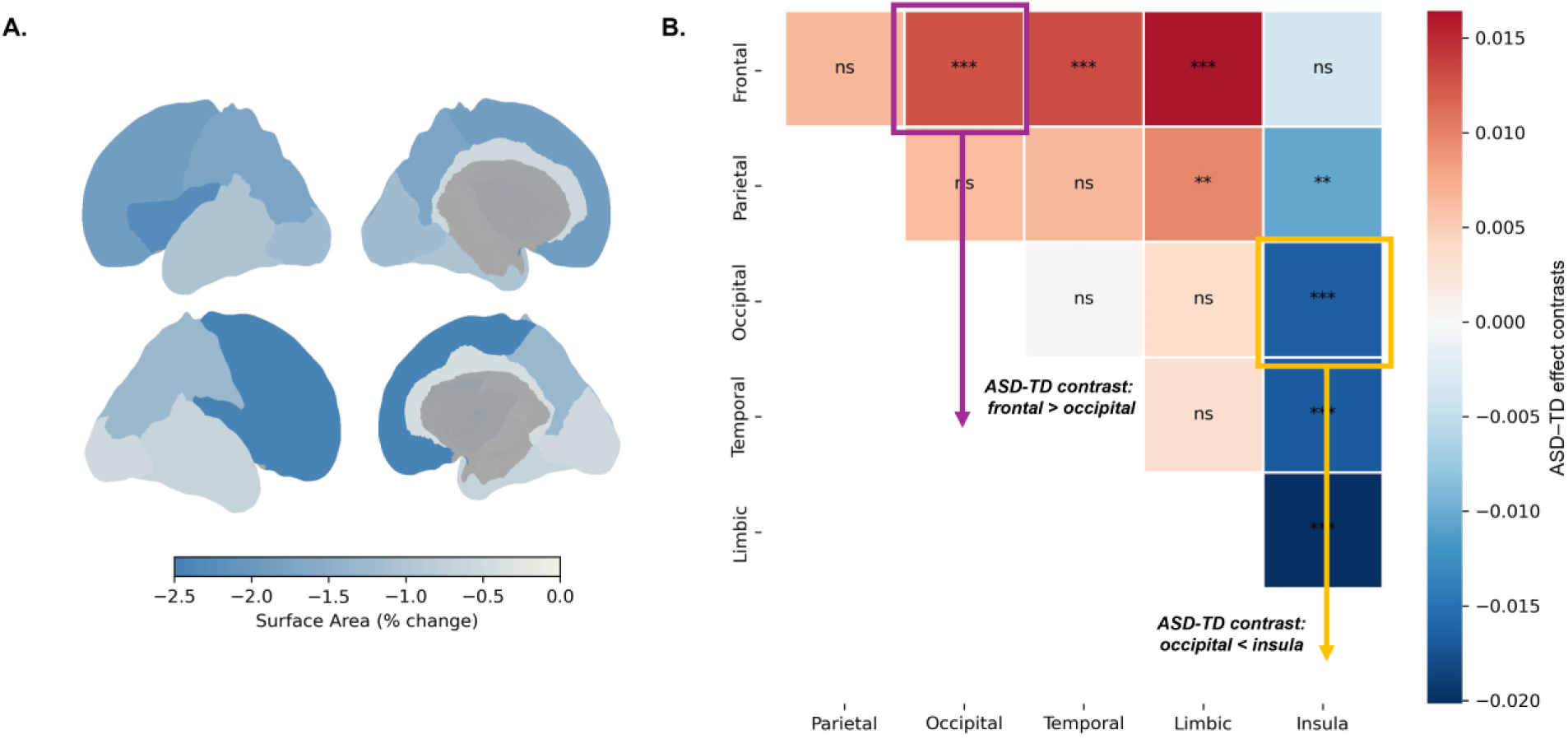
Relative ASD-TD differences summarizing results in SI-Table 9 (sensitivity analysis including acquisition quality mean QC and number of stacks as covariates). No relative changes are significant after accounting for data quality differences; the overall pattern of surface area reductions in ASD is maintained. **B.** Spatial heterogeneity in group effects from pairwise comparisons. Some lobes have significantly more pronounced surface area reductions in ASD-TD than others (row-column direction, i.e. red in row means given lobe is shows higher reductions than its pair in column, blue means less reduction). Significance codes: p<0.001 (***), p<0.01 (**), p<0.5 (*), p>0.5 (ns); FDR corrected.

**SI-Figure 7A.**
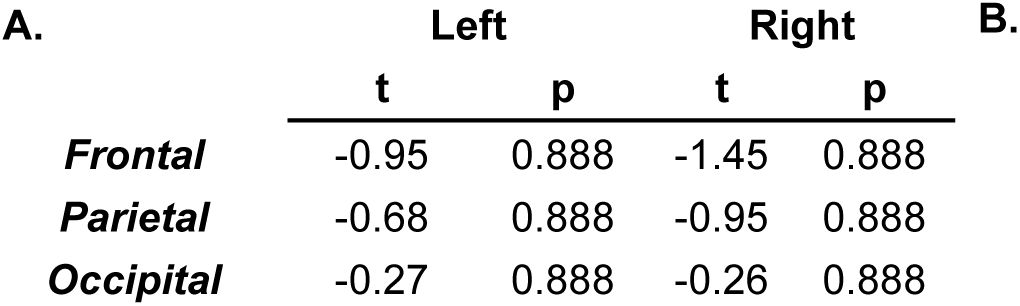
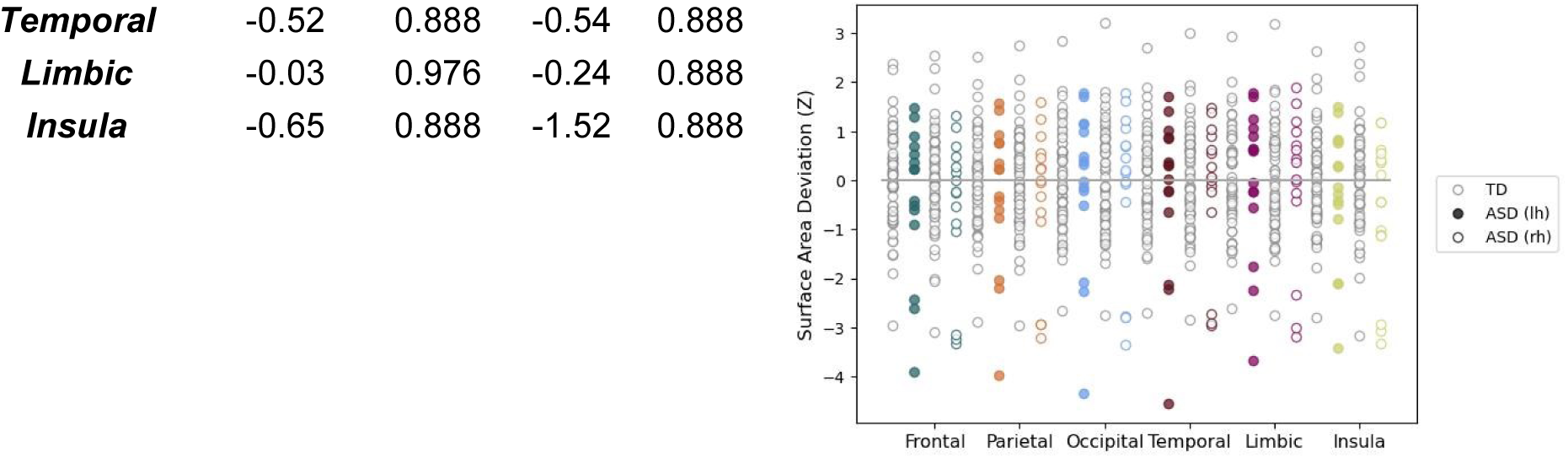
ASD normative deviations in lobar surface area (z-scores relative to TD normative model). Statistical significance of deviation from zero was assessed using mixed-effects estimated marginal means with FDR correction. Negative values indicate reduced surface area relative to TD expectations. (As z-scores are standardized relative to the TD normative model, mean Z values reflect effect sizes in units of TD standard deviations.) **B.** Scatter plots of subject-level deviation scores.

